# Miro1 expression alters global gene expression, ERK1/2 phosphorylation, oxidation, and cell cycle progression

**DOI:** 10.1101/2024.11.06.622334

**Authors:** Nathaniel Shannon, Cory Raymond, Chloe Palmer, David Seward, Brian Cunniff

## Abstract

Subcellular mitochondrial positioning in cells is necessary for localized energy and signaling requirements. Mitochondria are strategically trafficked throughout the cytoplasm via the actin cytoskeleton, microtubule motor proteins, and adaptor proteins. Miro1, an outer mitochondrial membrane adaptor protein, is necessary for attachment of mitochondria to microtubule motor proteins for trafficking. Previous work showed when Miro1 is deleted (Miro1^-/-^) from mouse embryonic fibroblasts (MEFs), the mitochondria become sequestered to the perinuclear space, disrupting subcellular energy and reactive oxygen species gradients. Here, we show that Miro1^-/-^ MEFs grow slower compared to Miro1^+/+^ and Miro1^-/-^ MEFs stably re-expressing the Myc-Miro1 plasmid. Miro1^-/-^ MEFs have a have a cell cycle defect with decreased percentage of cells in G1 and increased cells in the S phase of the cell cycle. We conducted the first ever RNA sequencing experiment dependent upon Miro1 expression and found differential expression in cell proliferation and migration genes upon deletion of Miro1, including the MAP Kinase signaling pathway. We find that ERK1/2 phosphorylation is elevated both spatially (cytoplasm and nucleus) and temporally following serum stimulation in Miro1^-/-^ MEFs. We investigated the expression levels and oxidation of the Dual Specificity Phosphatases (DUSP1-6), ERK1/2 target phosphatases. We found no differences in DUSP1-6 expression and oxidation under asynchronous and synchronized cells. Lastly, we evaluated the oxidation status of ERK1/2 and found an increase in ERK1/2 oxidation in the Miro1^-/-^ MEFs compared to Miro1^+/+^ and Myc-Miro1. These data highlight transcriptional control based off Miro1 expression and demonstrate the highly dynamic regulation of ERK1/2 upon deletion of Miro1 that may support the observed cell cycle and proliferation defects.

## Introduction

Subcellular mitochondrial positioning is critical for the cell to meet localized energy and signaling requirements. Mitochondria are strategically trafficked throughout the cytoplasm via the actin cytoskeleton, microtubule motor proteins, and adaptor proteins. Miro1, an outer mitochondrial membrane adaptor protein, is necessary for the attachment to microtubule motor proteins for mitochondrial trafficking [1–3]. Previous work from our lab and others shows when Miro1 is deleted (Miro1^-/-^) out of mouse embryonic fibroblasts (MEFs), the mitochondria become sequestered to the perinuclear space [4, 5]. This defect in subcellular mitochondrial positioning leads to increased reactive oxygen species (ROS) levels in the perinuclear/nuclear space and increased DNA damage. Conversely, peripheral ROS levels are significantly reduced in areas absent of mitochondria [4].

Mitochondrial positioning supports cell signaling cascades that govern diverse cellular phenotypes [5–16]. Hypoxia was shown to increase clustering of mitochondria around the nucleus in endothelial cells leading to an increase in ROS levels in the nucleus [17]. This supported oxidation of DNA bases in the hypoxia response element of the vascular endothelial growth factor (VEGF) promoter, leading to amplified binding of HIF1-α to the VEGF promoter and increased VEGF expression [17]. At the site of plasma membrane injury there is an influx of calcium which is lethal to the cell [18]. Mitochondrial trafficking and fragmentation at the plasma membrane was shown to be required to buffer the influx of calcium and prevent cell death [18]. The trafficked mitochondria are also necessary for local generation of ROS which is a key player in repairing the plasma membrane via redox signaling [19]. There is evidence indicating a role between peripheral mitochondria supporting cell migration. When AMP-activated protein kinase (AMPK) is activated, this causes mitochondria to traffic towards the leading edge of the cell which increases actin ruffling and cell migration. Inhibition of AMPK stops mitochondrial trafficking to the periphery which; therefore, prevents cell migration, indicating the importance of subcellular mitochondrial positioning [20]. In *C. elegans* the invasive anchor cell must break through the basement membrane during development. The invasive front of the anchor cell is enriched in the glucose transporters, FGT-1 and FGT-2, to increase intracellular glucose levels. The increased flux of glucose supports mitochondrial ATP production required for the reconstruction of the actin cytoskeleton supporting invasion [21, 22]. These examples provide evidence that mitochondrial subcellular positioning has diverse roles in numerous cell phenotypes.

Mitochondria also play a vital role in cell cycle progression. It was shown that mitochondria are more filamentous which supports increased ATP production during S phase [23]. In addition, mitochondrial-ER contact sites (MERCs) are expanded during mitosis which are essential for controlling the transfer of Ca^2+^ between the organelles. This is believed to aid in the production of ATP [24]. A decrease in ATP and increase in AMP, as well as reduced mitochondrial ROS (mROS), results in insufficient levels of cyclin E which is required for S-phase progression; therefore, stalling the cell in G1 [25].

The extracellular signal regulated kinase 1 and 2 (ERK1/2), members of the MAP-kinase signaling family, are involved in various signaling cascades during cell proliferation [26]. Similar to dynamic changes in mitochondrial structure and function during the cell cycle, ERK1/2 activity must be spatially and temporally regulated [26, 27]. In the G0-G1 transition, ERK1/2 must be dephosphorylated in the nucleus to allow a buildup of cyclin D1 to initiate the cell cycle [28]. In the G1 phase, ERK1/2 is phosphorylated (p-ERK1/2) and must remain phosphorylated to progress through G1 [29]. Progressing to S-phase, ERK1/2 must be phosphorylated and localized to the nucleus along with CDK2 so that cyclin E and CDK2 can form a complex to drive S-phase [30]. However, if the levels of p-ERK1/2 are too high this leads to an accumulation of p21^cip1^ (p21) which acts as a CDK inhibitor, thus leading to the disruption of the CDK2-cyclin E complex which is needed for progression of the S-phase [31]. At the end of the S-phase, p-ERK1/2 levels are high, while at the end of mitosis p-ERK1/2 is rapidly dephosphorylated to allow exit from the cell cycle [32]. ERK1/2 activity can be altered in two primary ways: first, by dephosphorylation via phosphatases, primarily the dual-specificity phosphatases (DUSPs) [33]. Secondly, by ERK1/2 oxidation which regulates its phosphorylation [34]. The activity of DUSPs is negatively regulated by oxidation of the active site, leading to prolonged ERK1/2 phosphorylation [34, 35]. ERK1/2 itself can become oxidized, causing persistent phosphorylation and sustained signaling responses [36]. Specifically, ERK2 is vulnerable to direct oxidation on Cys159 which is located in its D-recruitment site binding domain [37]. Therefore, the regulation of ERK1/2 in space and time is necessary for the progression of the cell cycle.

In this study we found that deletion of Miro1 leads to reduced cell proliferation. We conducted the first ever RNA sequencing experiment dependent upon Miro1 expression and found differentially expressed genes associated with cell migration and cell proliferation, including the MAPK pathway. We found that ERK1/2 is hyper-phosphorylated in Miro1^-/-^ MEFs, both spatially (cytoplasm and nucleus) and temporally (following serum stimulation). We further investigated the phosphatases responsible for the dephosphorylation of ERK1/2 by looking at their expression levels as well as oxidation state. We found no difference in the expression and oxidation of the ERK1/2 DUSPs. Lastly, we show that ERK1/2 oxidation is increased in the Miro1^-/-^ MEFs compared to Miro1^+/+^ and Myc-Miro1 following serum stimulation. Together, these studies offer new insights into the role of Miro1 protein expression in regulating gene-expression, cell cycle progression and redox dependent signaling.

## Results

### Deletion of Miro1 leads to perinuclear mitochondrial clustering, increased networking, and no change to matrix H2O2 levels

Miro1 deletion leads to mitochondrial clustering around the nucleus in numerous cell types [1, 4–6, 9, 38]. Utilizing wild-type (Miro1^+/+^) mouse embryonic fibroblasts (MEFs), Miro1 knockout MEFs (Miro1^-/-^) and a Miro1^-/-^ stable cell line stably transfected with a Myc-Miro1 DNA plasmid (referred to as Myc-Miro1), we conducted immunofluorescence staining with an antibody for Translocase of the Outer Mitochondrial Membrane 20 protein (TOMM20) to confirm these results (**Fig. 1A**). Miro1^+/+^ and the Myc-Miro1 rescue cell line show similar mitochondrial distribution throughout the cell while Miro1^-/-^ MEFs show a ∼40% decrease in mitochondrial occupancy of the cell periphery (**Fig. 1B**). TOMM20 fluorescence intensity was decreased in the Myc-Miro1 immunofluorescence staining, however Western blotting of cell lysates showed no significant difference in TOMM20 levels among the three cell types (**Fig. 1 C-E**). Previous work in yeast and mammalian cells has shown that Miro1 may play a role in structural transitions of mitochondria [39–41]. Form factor analysis, a measurement of mitochondrial networking, shows a significant increase in mitochondrial networking in Miro1^-/-^ MEFs (**Fig. 1F**), a phenotype previously observed by others [40].

**Figure 1:**
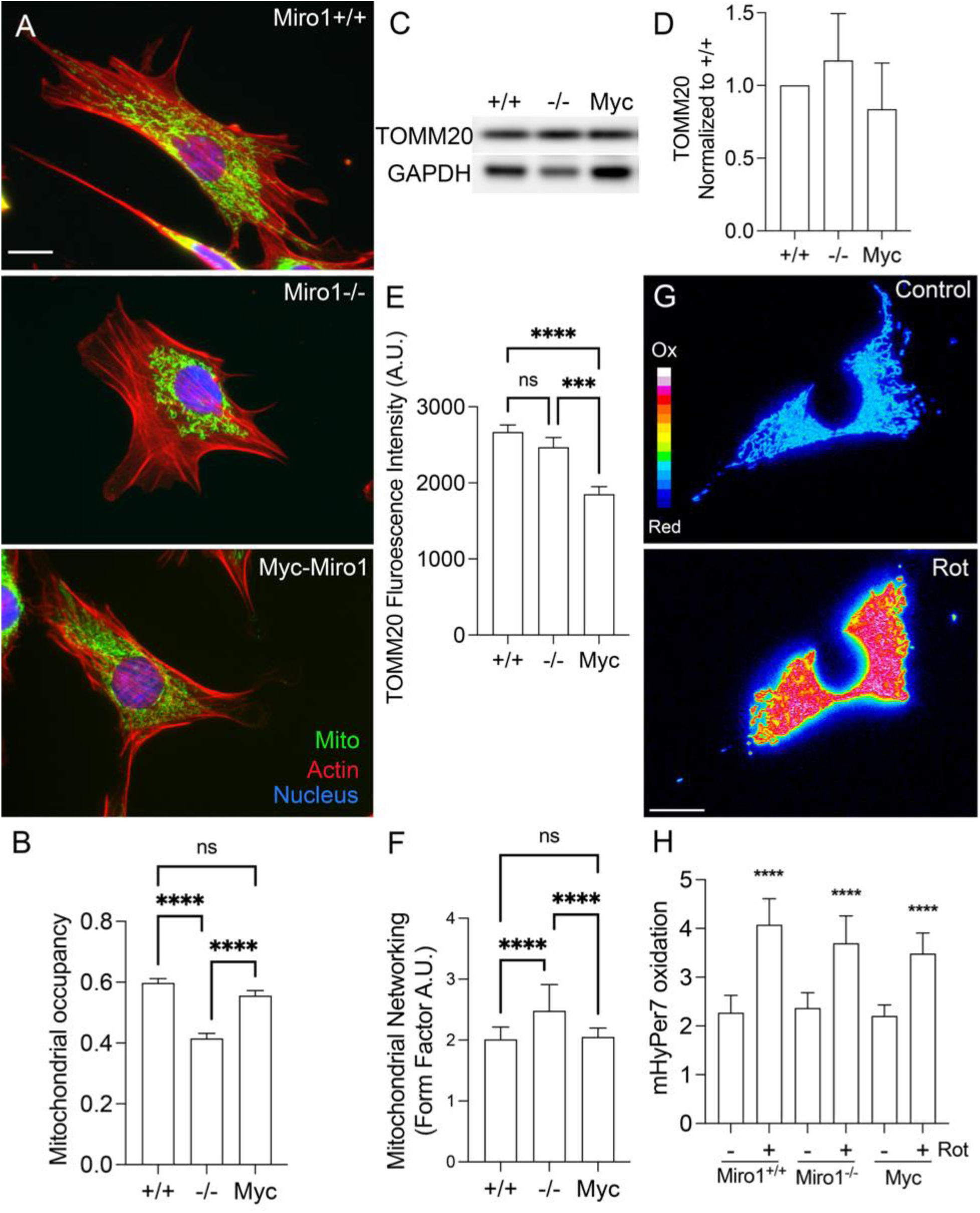
Deletion of Miro1 leads to perinuclear mitochondrial clustering, increased networking without changes to matrix H2O2 levels. **A**) Mitochondria occupy the peripheral and perinuclear space in wild-type mouse embryonic fibroblasts (top). Miro1 deletion (Miro1^-/-^) results in mitochondrial perinuclear clustering (middle). Rescue of Miro1 via Myc-Miro1 expression recovers the wild-type mitochondrial phenotype (bottom). B) Quantification of mitochondrial occupancy in the cell (n=50 cells). C – D) Western blot of TOMM20 and GAPDH with densitometry quantification (n=3 biological replicates). E) Mean fluorescence intensity of the TOMM20 staining (n=26 cells). F) Quantification of mitochondrial networking via form factor analysis (n=20 cells). G) (Top) Representative image of Miro1^+/+^ MEF transfected with MLS-HyPer7, (Bottom) Cell treated with 5 µM rotenone (Rot) for 40 minutes. H) Quantification of ratiometric HyPer7 value pre and post rotenone treatment (n=18 cells). *** p < 0.001, **** p < 0.0001, SEM shown, One-Way ANOVA with Tukey’s posttest.

Previous studies using the genetically encoded cytoplasmic/nuclear expressed H2O2-sensitive fluorescent biosensor HyPer7 showed a decrease in HyPer7 oxidation in the cell periphery devoid of mitochondria in Miro1^-/-^ cells and an increase at perinuclear and nuclear areas [4]. HyPer7 is pH stable and has two excitation peaks at 400 and 499 nm that have variable excitation profiles dependent on oxidation of the probe [42]. A higher numerical ratio reflects a higher amount of oxidized HyPer7 probe, and therefore a higher local level of H2O2. To confirm that previously noted differences in intracellular oxidation are due to mitochondrial localization and not the levels of H2O2 being produced within the mitochondria, we utilized a matrix localized (MLS) HyPer7 probe [42]. Miro1^+/+^, Miro1^-/-^ and Myc-Miro1 MEFs transfected with MLS-HyPer7 probe were visualized by live cell imaging to determine mitochondrial H2O2 levels. 5µM of Rotenone, a mitochondrial complex I inhibitor, was added after 10 minutes as a positive control (**Fig. 1G #x0026; H**). MLS-HyPer7 oxidation was consistent between all three cell types, demonstrating that H2O2 levels within the mitochondrial matrix are the same. This suggests differences in HyPer7 oxidation in the cytoplasm and nucleus are likely due to mitochondrial localization [4], not the differences in the amount of H2O2 being produced between cell types. Together, these results indicate loss of Miro1 in MEFs leads to perinuclear clustering of mitochondria, increased mitochondrial networking with no change in mitochondrial matrix levels of H2O2.

### Global gene expression changes associated with Miro1 loss

Cell and animal studies have uncovered important processes associated with the expression of Miro1 [16, 43]. To globally assess gene expression changes dependent on Miro1 expression we conducted RNA sequencing of Miro1^+/+^, Miro1^-/-^ and Myc-Miro1 MEFs. PCA analysis shows strong clustering of biological replicates from each cell line (n = 4, **Supplemental Fig. 1**). We identified 1897 genes with > 2-fold difference in expression with an adjusted p-value <0.05 in Miro1^-/-^ MEFs compared to Miro1^+/+^ MEFs (**Fig. 2A and Supplemental Table 1).** We focused our analysis of this data on genes that are differentially expressed in Miro1^-/-^ MEFs (KO) compared to Miro1^+/+^ (WT) and rescued by Myc-Miro1 re-expression. An example is the Cdh2 (n-cadherin) gene which is significantly downregulated in Miro1^-/-^ MEFs relative to WT and rescued in Myc-Miro1 MEFs (**Fig. 2B**). Cdh2 expression was validated by RT-qPCR (**Supplemental Fig. 1**). Deletion of Miro1 leads to a cell migration defect [5] and we find numerous genes associated with cell migration and attachment to be downregulated in Miro1^-/-^ MEFs compared with WT (eg. *Ezr*, *Cdh2, Cadm1, Pkp1, Edil3*) (**Fig. 2A & B, Supplemental Fig. 1 and Supplemental Table 1**). Perinuclear clustering of mitochondria following hypoxia has been shown to support HIF-1α dependent transcription of the Vegfr gene [17]. We find a significant increase in both Vegfr1 (*Flt1*) and Vegfr2 (*Kdr*) expression in Miro1^-/-^ MEFs with perinuclear clustered mitochondria and these expression changes are ameliorated by Myc-Miro1 re-expression (**Fig. 2A & B, Supplemental Fig. 1 and Supplemental Table 1)**. We also observed significant changes in the expression of genes associated with cell growth and response to growth factors including the Pdgf and Egf receptors (*Pdgfra and Egfr)*) (**Fig. 2A & B, Supplemental Fig. 1 and Supplemental Table 1**). To identify candidate biologic processes impacted by Miro1 deletion we next queried our expression data using KEGG analysis. This approach identified significant enrichment in several pathways and processes including Ras, ErbB, Focal Adhesion, VEGF, HIF-1, and MAPK signaling (**Fig. 2C and Supplemental Table 2).** Together these data show significant changes in Miro1 dependent gene expression, implying a critical role for mitochondrial localization on regulating cellular transcriptional patterns. We further evaluated the MAPK signaling pathway in subsequent experiments.

**Figure 2:**
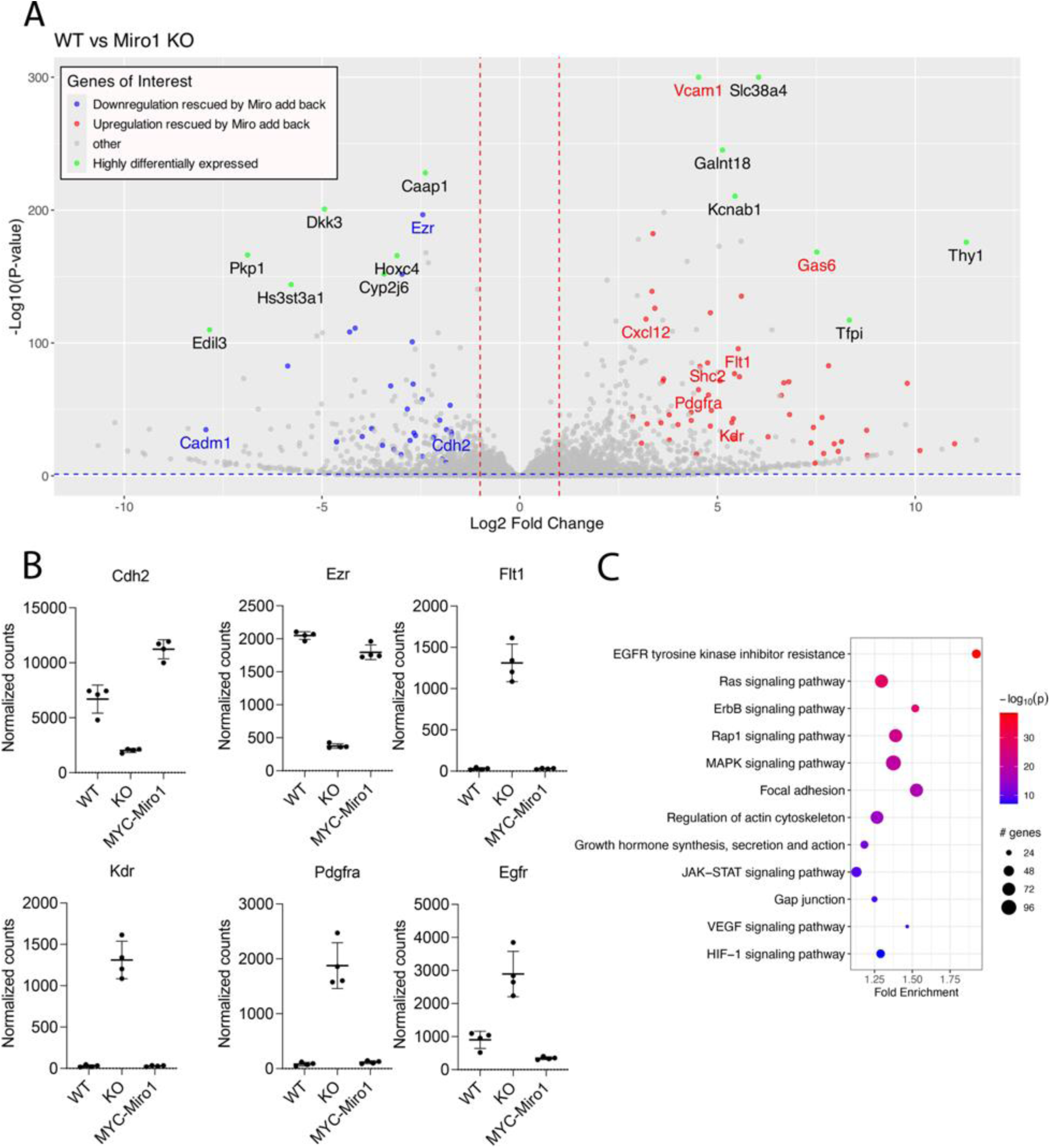
Global gene expression changes associated with Miro1 loss. A) Volcano plot of differential gene expression between Miro1^+/+^, Miro1^-/-^ and Myc-Miro1. B) Examples of genes with differential gene expression between Miro1^+/+^ (WT), Miro1^-/-^ (KO) that are rescued by Myc-Miro1 expression. C) KEGG analysis of differentially expressed genes.

### Miro1^-/-^ MEFs proliferate slower and have a cell cycle defect at G1 and S phase

While passaging cells we observed that Miro1^-/-^ MEFs grew slower than Miro1^+/+^ or Myc-Miro1 MEFs. To quantify this observation, we conducted cell proliferation assays. Miro1^-/-^ MEFs grew slower compared to that of Miro1^+/+^ as well as Myc-Miro1 beyond 72 hours of growth (**Fig. 3A**). All the cell lines were viable throughout the 144 hours. To further investigate cell proliferation and cell cycle progression, we conducted propidium iodide staining and flow cytometry. Miro1^-/-^ MEFs showed a significant decrease in the population of cells in the G0/G1 cell cycle stage and an increase in the population of cells in the S cell cycle stage (**Fig. 3B & C**). Miro1^+/+^ and Myc-Miro1 MEFs had an expected normal cell cycle distribution. We next evaluated cyclin A-E baseline expression via Western blotting because these proteins are temporally expressed throughout different cell cycle stages [44]. Miro1^-/-^ MEFs had a ∼40% increase in cyclin D1 expression, a ∼40% increase in cyclin A2 expression and a ∼300% increase in cyclin B1 expression compared to Miro1^+/+^ and Myc-Miro1 MEFs (**Fig. 3D & E**). We attempted to look at cyclin E expression but were unsuccessful with the use of multiple antibodies. These data suggests that deletion of Miro1 leads to a cell cycle and proliferation defect associated with changes in cyclin expression.

**Figure 3.**
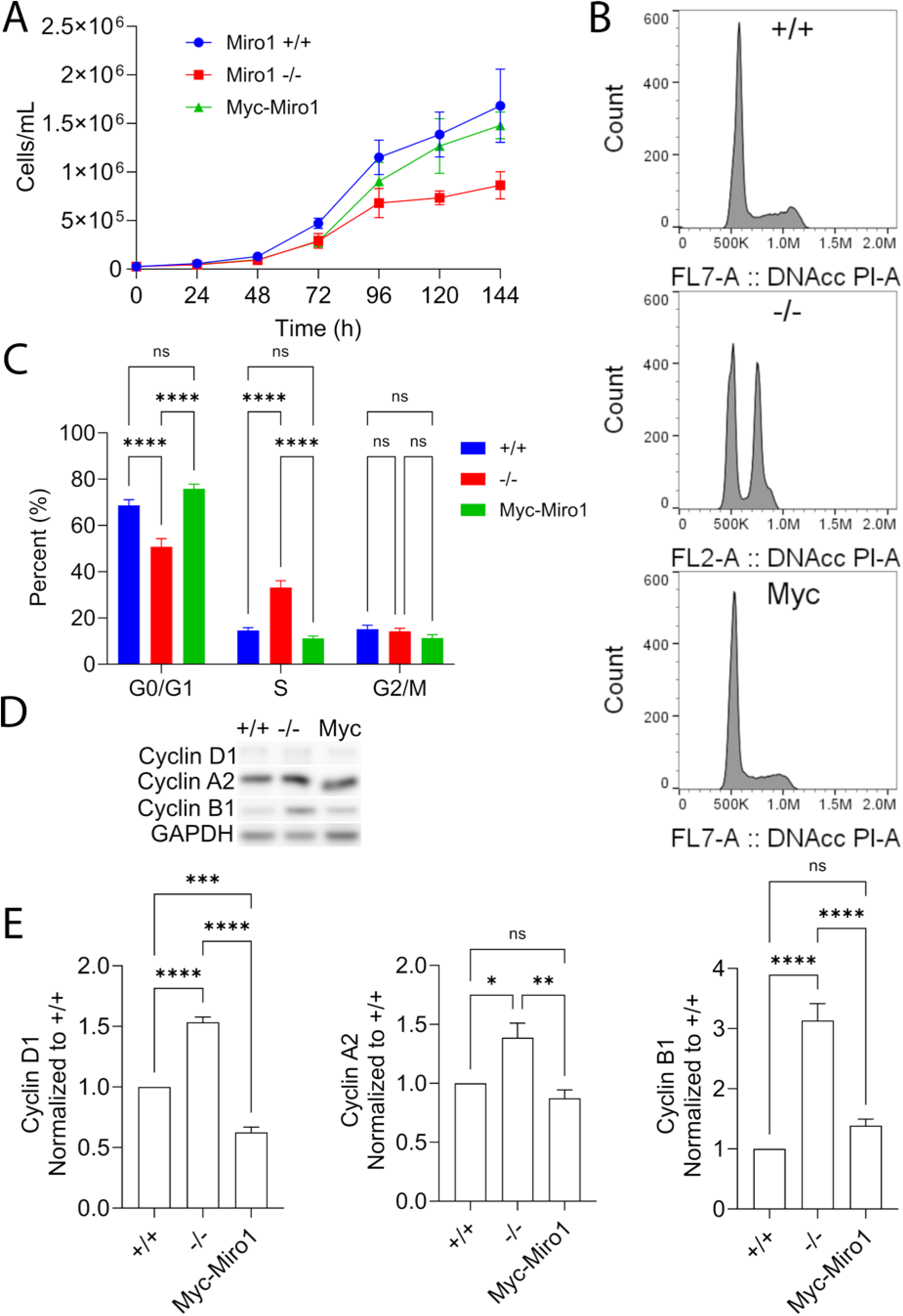
Miro1^-/-^ MEFs display slower proliferation as well as a stall in the G2/M transition of the cell cycle. A) Cell proliferation of Miro1^+/+^, Miro1^-/-^, and Myc-Miro1 MEFs measured every 24 hours for 144 hours (n=3 biological replicates). B) Propidium iodide staining of Miro1^+/+^, Miro1^-/-^, and Myc-Miro1 MEFs measured by flow cytometry (n=4-6 biological replicates). C) Quantification of each stage of the cell cycle using the FlowJo software accompanied by Watson pragmatic algorithm (Two-way ANOVA with Tukey’s posttest). D) Whole cell lysates of asynchronous cells analyzed via Western blots and probed for the G2-phase and mitotic cyclins, cyclin A2 and cyclin B1 respectively. E) Quantification of the Western blots in C normalized to GAPDH (n=4 biological replicates). * p < 0.05, ** p < 0.01, **** p < 0.0001, SEM shown, One-way ANOVA with Tukey’s posttest.

### ERK1/2 phosphorylation is increased temporally and spatially in Miro1^-/-^ MEFs

We have shown that Miro1^-/-^ MEFs have a cell cycle and proliferation defect and dysregulation in genes involved in the MAPK pathway (**Fig. 1 - 3**). The MAPK pathway plays a critical role in cell proliferation and cell cycle progression [26]. Therefore, we investigated MAPK pathway proteins for activation via phosphorylation. Phosphorylation of mitogen-activated protein kinase kinase (MEK1/2), the kinase upstream of ERK1/2, which is activated following growth factor addition was similar between cell lines (**Fig. 4A & C**). We next looked at the substrate for MEK1/2, ERK1/2 and found that ERK1/2 is hyperphosphorylated in Miro1^-/-^ MEFs compared to Miro1^+/+^ and Myc-Miro1 (**Fig. 4B & D**). ERK1/2 is trafficked between the cytoplasm and nucleus for co-transcriptional activity. We conducted cell fractionization and found ERK1/2 is hyperphosphorylated in Miro1^-/-^ MEFs in both compartments (**Fig. 4E & F**). Additionally, cyclin B1, the G2 cyclin, was elevated in the nuclear fraction of the Miro1^-/-^ MEFs (**Fig. 4E & G**), corroborating our observation that Miro1^-/-^ MEFs are preferentially in the S/G2M cell cycle phase (**Fig. 3**).

**Figure 4:**
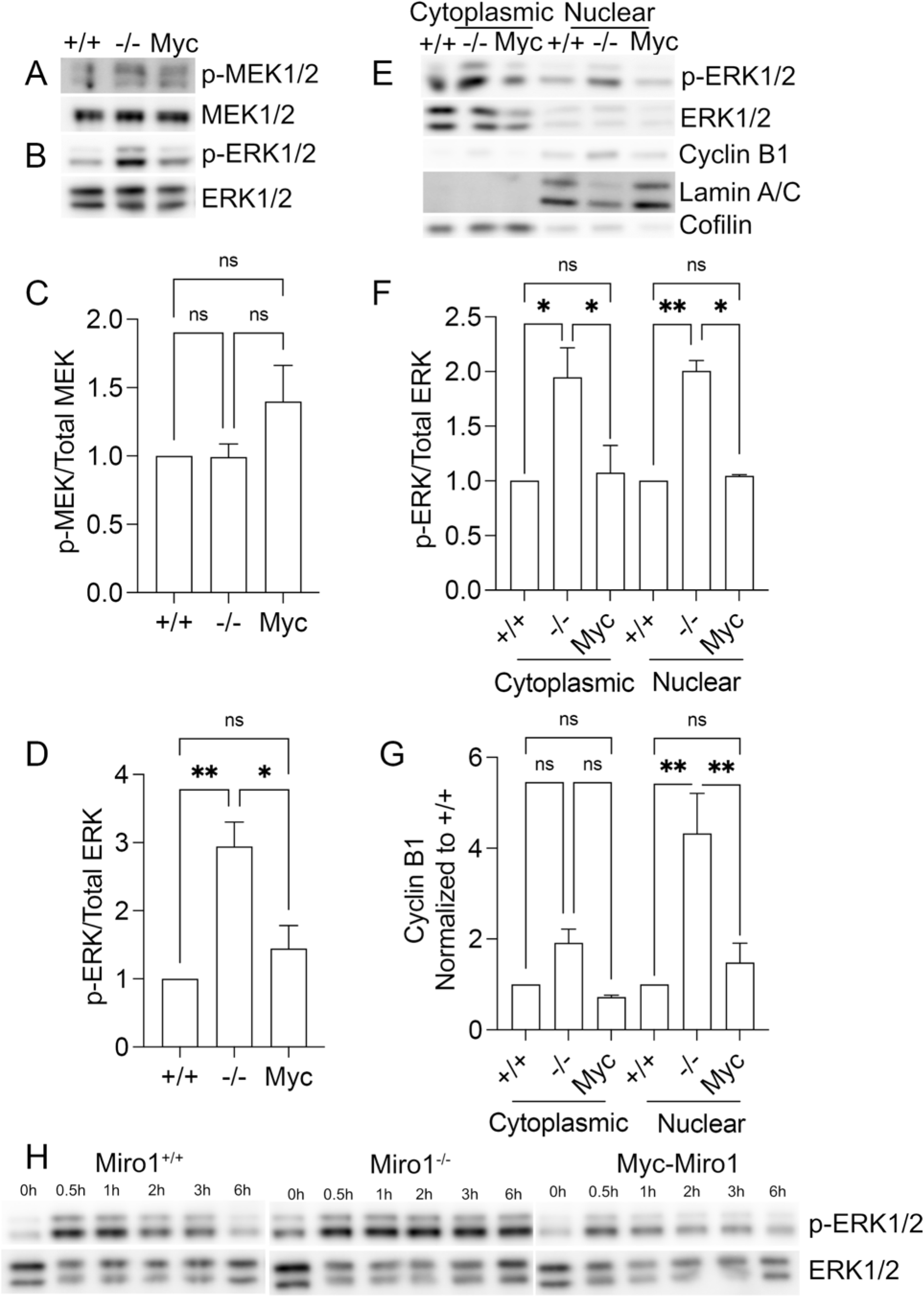
ERK1/2 phosphorylation is increased temporally and spatially in Miro1^-/-^ MEFs. A and B) Whole cell lysates of asynchronous cells were analyzed via Western blots and probed for phosphorylated and total proteins in the MAPK pathway (n=3 biological replicates). C and D) Quantification of A and B, respectively (n=3 biological replicates). E) Western blots of phospho-ERK1/2, total ERK1/2, and cyclin B1 in the cytoplasmic and nuclear fractions. Lamin A/C was used for the nuclear loading control and cofilin was used as the cytoplasmic loading control (n=3 biological replicates). F) Densitometry of phospho-ERK1/2 normalized to total ERK1/2 from nuclear and cytoplasmic fractions (n=3 biological replicates). G) Densitometry of cyclin B1 in the cytoplasmic and nuclear fractions (n=3 biological replicates). H) Phospho-ERK1/2 and total ERK1/2 Western blots from cells starved for 24 hours in 1% FBS followed by full serum addition (10%) for the indicated time points. Lysates were generated at the indicated time points (n=4 biological replicates). * p < 0.05, ** p < 0.01, SEM shown, One-way ANOVA with Tukey’s posttest.

ERK1/2 is dynamically regulated throughout the cell cycle by phosphorylation and dephosphorylation [45], therefore we evaluated the temporal phosphorylation of ERK1/2 following serum starvation and stimulation in our MEF cell lines. Cells were starved in 1% FBS for 24 hours then full serum was added (10% FBS) for time points between 0.5 hours and 6 hours. We determined the temporal phosphorylation of ERK1/2 via Western blot (**Fig. 4H**). Baseline phosphorylation of ERK1/2 under serum starvation was slightly elevated in Miro1^-/-^ compared to Miro1^+/+^ and Myc-Miro1 MEFs (**Fig. 4H**). Upon serum addition in all three MEF cell lines, there was rapid phosphorylation of ERK1/2 which peaked at 0.5 hours for all three MEF cell lines (**Fig. 4H**). In Miro1^+/+^ and Myc-Miro1 MEFs ERK1/2 was slowly dephosphorylated throughout the remaining time points until it approached baseline phosphorylation levels at 6 hours (**Fig. 4H**). In the Miro1^-/-^ MEFs, ERK1/2 phosphorylation remained elevated throughout the 6-hour time point. These data together suggest a dysregulation in the dynamics of ERK1/2 in Miro1^-/-^ MEFs following serum starvation and stimulation.

### The ERK1/2 dual specificity phosphatases (DUSPs) are unchanged in their expression and oxidation

DUSPs and PP2AC regulate MAPK activity via dephosphorylation which renders the kinase inactive [33, 46]. Oxidation of phosphatases, specifically DUSPs, renders them inactive; therefore, they are unable to dephosphorylate their target kinase, resulting in the kinase remaining phosphorylated [34, 35]. Our data shows increased perinuclear H2O2 levels in Miro1^-/-^ MEFs [4] and increased ERK1/2 phosphorylation (**Fig. 4**), therefore we hypothesized that the DUSPs may be dysregulated, via expression and/or oxidation in the Miro1^-/-^ MEFs. Expression of DUSP1/3/5/6 and PP2AC was similar across all three cell lines (**Fig. 5A & B**). We next labeled oxidized proteins in the cell lysates using DCP-Bio1, a cell permeable version of biotin labeled dimedone which will covalently adduct protein sulfenic acids [47]. Labeled proteins were immunoprecipitated using streptavidin beads and protein oxidation was analyzed by Western blots. DUSPs and PP2AC oxidation did not vary between the MEF cell lines (**Fig. 5A & C**).

**Figure 5:**
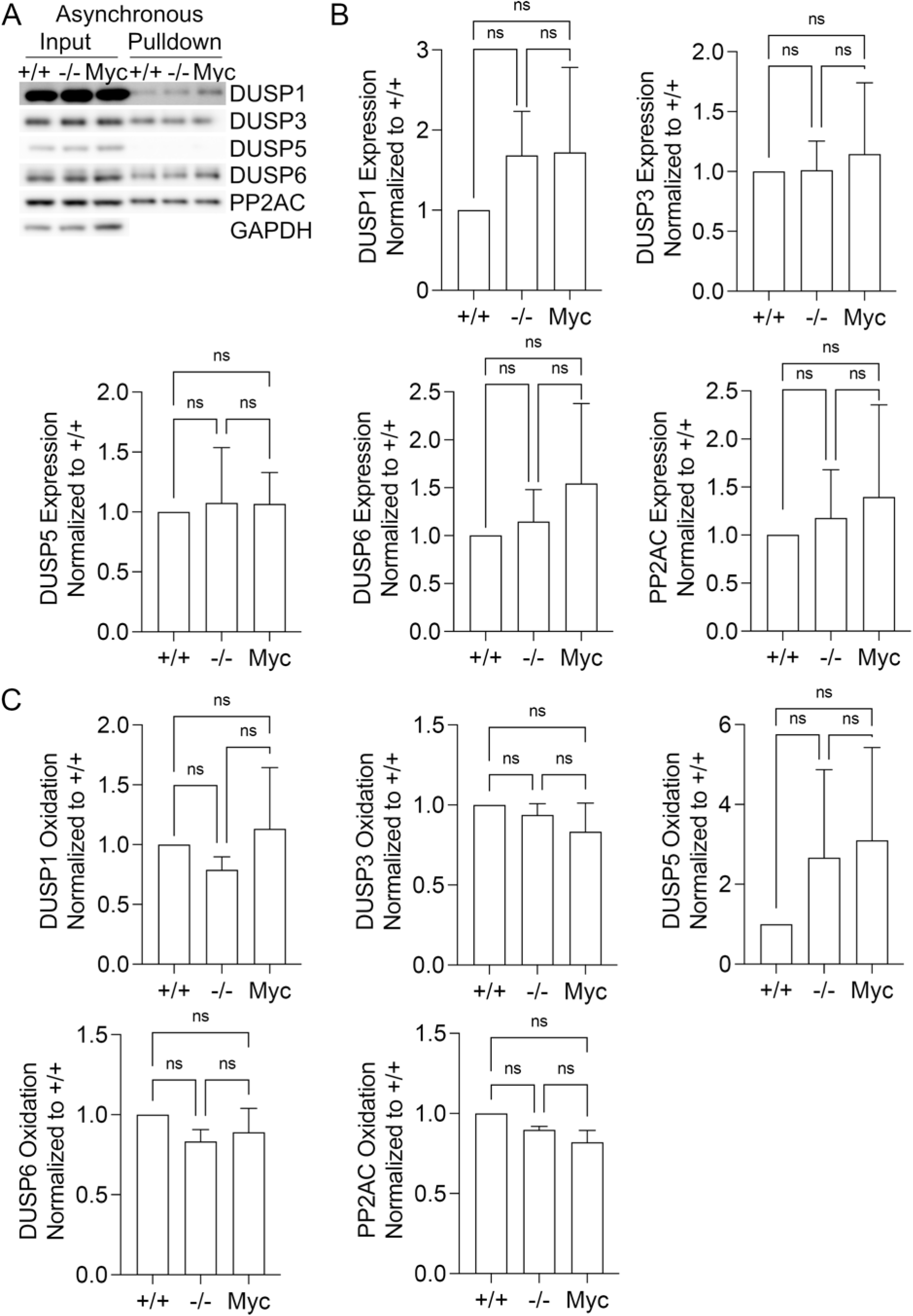
The expression of the ERK1/2 phosphatases, dual specificity phosphatases (DUSPs), is unchanged in Miro1^-/-^ MEFs. A) Asynchronous cells were treated with DCP-Bio1 and the following DUSPs were analyzed and quantified for relative abundance of oxidation. B) Relative abundance of each DUSP was quantified via the whole cell lysate input, in A, compared to GAPDH then normalized to Miro1+/+. C) Bar graph quantifying the pulldown of the oxidized protein compared to the whole cell lysate input then normalized to Miro1+/+. (n=3 biological replicates, One-way ANOVA with Tukey’s posttest).

Our cell cycle data indicates Miro1^-/-^ MEFs are disproportionally present in the S phase of the cell cycle (**Fig. 3B & C**). To evaluate DUSP expression and oxidation specifically at S and G2/M cell cycle phases we serum starved the cells in 0.1% FBS for 48 hours then added back serum (10% FBS) for the indicated time points and collected lysates (**Fig. 6 A & B**). We monitored the expression of the cyclin proteins by protein Western blotting (**Fig. 6 A & B**). Cyclin D1 is rapidly expressed following serum stimulation and peaked at ∼12 hours equally in all three cell lines (**Fig. 6B**). We failed to evaluate cyclin E expression due to poor antibodies. Cyclin A2 expression was increased at 18-20 hours before returning to baseline around 24 hours in all cell lines (**Fig. 6B**). Cyclin B1 expression was elevated in serum starved Miro1^-/-^ MEFs and was increased at all time points compared to Miro1^+/+^ and Myc-Miro1 MEFs (**Fig. 6B**). Cyclin B1 expression peaked at 20 hours (**Fig. 6B**), in line with results in **Fig. 3D & E** indicating increased cyclin B1 expression in asynchronized cells. We next evaluated the expression and oxidation of the DUSPs at 20 hours following serum stimulation (**Fig. 6C & D**). We were unable to detect DUSP5 and PP2AC in the synchronized cells. We observed no change in DUSP oxidation in cells following serum stimulation at a time when cyclin A2 and cyclin B1 expression is maximal.

**Figure 6.**
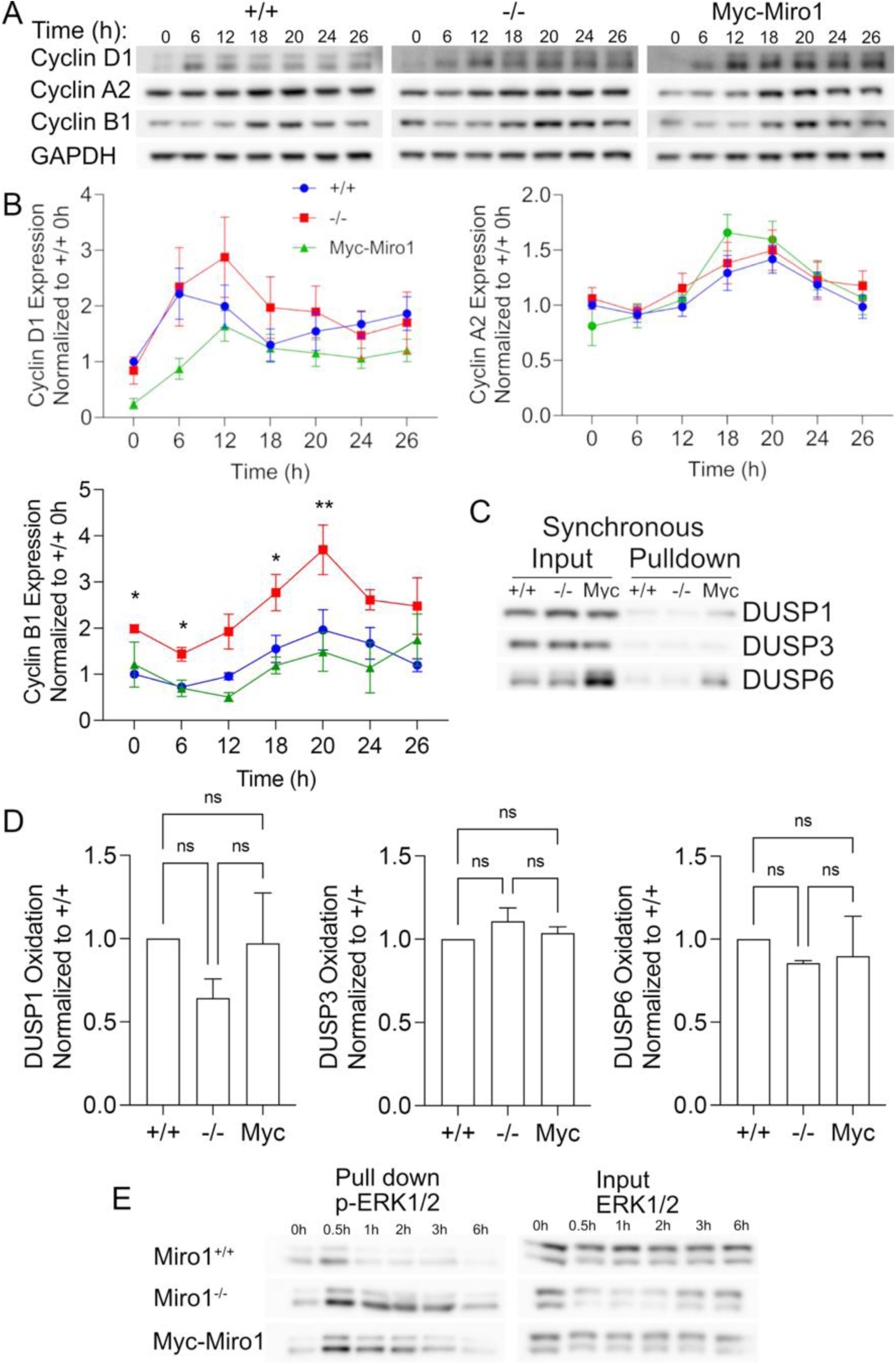
DUSP expression and oxidation is equal in MEF cell lines at S/G2M while ERK1/2 oxidation is prolonged in Miro1-/-MEFs. A) Cells were serum starved (0.1% FBS) for 48 hours then treated with full serum (10% FBS) for the indicated time points. The whole cell lysates were analyzed via Western blots for the presence of the cyclins that are expressed at various stages of the cell cycle. B) Quantification of (A) of the expression of cyclin D1, A2, and B1 post serum starvation (n=4 biological replicates). C) Cells were serum starved (0.1% FBS) for 48 hours then released in full serum (10% FBS) for 20 hours and then collected in the presence of DCP-Bio1. The following DUSPs were analyzed via Western blots. D) Quantification of DUSP oxidation via pulldown divided by the input and normalized to Miro1^+/+^ (n=4 biological replicates). E) Cells were serum starved (1% FBS) for 24 hours then treated with full serum (10% FBS) for the indicated time points. Cells were treated with DCP-Bio1 and analyzed via Western blot for total ERK1/2 on the whole cell lysate portion and for p-ERK1/2 on the pulldown section. * p < 0.05, ** p < 0.01, SEM shown, One-way ANOVA with Tukey’s posttest.

### ERK 1/2 oxidation is prolonged in Miro1^-/-^ MEFs following serum stimulation

The oxidation of ERK1/2 has been described to occur in response to proliferative signals [36], oxidative stress [37] and by direct bolus addition of H2O2 or nitric oxide donors to cell culture [48]. Oxidation of ERK1/2 strongly modulates ERK1/2 kinase activity *in vitro* and substrate selection for ERK2 [37]. Therefore, we evaluated ERK1/2 oxidation following serum stimulation using DCP-Bio1 labeling and immunoprecipitation. ERK1/2 is rapidly oxidized following serum stimulation in all three cell lines (**Fig.6E**). In Miro1^-/-^ MEFs oxidation of ERK1/2 is sustained past 2-4 hours compared to Miro1^+/+^ MEFs (**Fig. 6E**). These data show that the temporal oxidation status of phosphorylated ERK1/2 is prolonged in Miro1^-/-^ MEFs. Together, ERK phosphorylation and oxidation dynamics are dysregulated in Miro1-/-MEFs and correlate with changes in cell cycle protein expression and dynamics.

## Discussion

In this study we provide evidence that Miro1 expression supports mitochondrial trafficking, cell proliferation, cell cycle dynamics and the expression of key genes associated with cell migration, proliferation and growth signaling. Deletion of Miro1 leads to elevated and prolonged ERK1/2 phosphorylation and oxidation following serum stimulation. Our findings uncover novel features of Miro1 expression associated with cell proliferation and provide a robust RNA-sequencing data set for future investigation of molecular mechanisms governed by Miro1 expression and mitochondrial dynamics.

Deletion of Miro1 and 2 in HEK293 and U2OS cells was shown to lead to asymmetric mitochondrial distribution to daughter cells by disrupted interaction between Miro and CENPF on the growing tips of microtubules, supporting a cell cycle defect [13]. Our data indicates an earlier defect in cell cycle progression as asynchronous Miro1^-/-^ MEFs are disproportionately present in S phase with a decrease in cells in G1 (**Fig. 3B & C**). This is supported by aberrant expression of cyclin D1, cyclin A2 and cyclin B1 in asynchronous Miro1^-/-^ MEFs. As cyclins must be temporally and spatially expressed to support cyclin dependent kinase (CDK) activity at specific cell cycle phases [44], the mistimed or dysregulation of expression of cyclins may underly the observed cell cycle defects. In addition, we find increased cyclin B1 expression in the nucleus which supports interaction with CDK1. Aberrant CDK1 activity has been shown to induce S phase failure in MEFs by inducing premature mitotic events during S phase [49]. We also observed increased mitochondrial networking in Miro1-/-MEFs which needs further investigation due to the dynamic nature of mitochondria throughout the cell cycle. Together, our data supports the mis-regulation of cyclin expression that correlates with the observed cell cycle defect and previously published results [50].

We provide for the first time an RNA-sequencing data set evaluating differentially expressed genes dependent on Miro1 expression in MEFs. Pathways of great interest and previously shown to be influenced by either Miro1 expression or perinuclear mitochondrial clustering are identified. These include cell migration genes (CDH2, CDH13, EZR), response to growth factor signaling (PDGFRA, EGFR) and angiogenesis (FLT, KDR). Additional KEGG analysis highlights pathways of significant interest that are represented by differentially expressed genes in Miro1-/-MEFs. These pathways and the genes associated with them will be the basis of additional studies.

Miro1 is a critical mitochondrial adapter protein required for subcellular trafficking of mitochondria. Additionally, Miro1 is important for peroxisome trafficking and biogenesis, tethering of mitochondria to the endoplasmic reticulum and mitophagy [7, 8, 11, 51]. Therefore, our MEF cell lines in which we rescue Miro1 deletion through stable expression Myc-Miro1 provides an optimal system to evaluate changes dependent on Miro1 expression. By deleting out Miro1 in MEFs we see that there is a defect in the regulation of ERK1/2 phosphorylation and not due to the upstream kinase, MEK1/2, which had no change in its phosphorylation status. ERK1/2 phosphorylation levels are similar in Miro1^+/+^ and Myc-Miro1 MEFs indicating that Miro1 expression is important in the regulation of its phosphorylation. This could be due to perinuclear clustering of mitochondria or other Miro1 functions. We investigated ERK1/2 localization and dynamics due to its involvement in cell cycle regulation and proliferation. ERK1/2 needs to be phosphorylated and localized in the nucleus to elicit its responses on the cell cycle [52, 53], but also needs to be de-phosphorylated towards the end of mitosis to allow the cell to exit the cell cycle [32]. We observed that ERK1/2 phosphorylation is elevated in the cytoplasmic fraction as well as the nuclear fraction of Miro1^-/-^ MEFs; therefore, ERK1/2 is able to localize properly. Similarly, cyclin B1 expression is increased in the nucleus of Miro1^-/-^ MEFs which is associated with causing premature mitotic events in the S phase [49]. The localization of ERK1/2 is proper for eliciting its responses on the cell cycle. Upon the addition of serum, ERK1/2 phosphorylation remains elevated and is not dephosphorylated in a time dependent manner in the Miro1^-/-^ MEFs, whereas we see progressive dephosphorylation in a time dependent manner in the Miro1^+/+^ and Myc-Miro1 MEFs. Therefore, its dephosphorylation is disrupted whcih led us to look at phosphatases governing ERK1/2 phosphorylation.

First, we looked at the baseline expression of DUSPs, ERK1/2 phosphatases, and found no change under asynchronous conditions. It is known that phosphatases can be deactivated upon oxidation [54] and our previous studies show that Miro1^-/-^ MEFs have an oxidant rich perinuclear and nuclear area [4]. This led us to question the oxidation status of these phosphatases given the spatial localization and hyperphosphorylation of ERK1/2 in the nucleus of the Miro1^-/-^ MEFs. We labeled the oxidized proteins using DCP-Bio1 and observed no differences in the protein oxidation of DUSP1,3, 5 and 6 as well as PP2AC. Because there is increased cells in the S phase in the Miro1^-/-^ MEFs, we synchronized the cells in S/G2M. We chose to look at S/G2M because upon synchronization we do not observe a difference in the expression of cyclin A2, but rather see an increase in the expression of cyclin B1 in the Miro1^-/-^ MEFs. Overall, we do not observe any differences in expression or oxidation of the DUSPs after the cells were synchronized in mitosis. We attempted to look at DUSP5 and PP2AC oxidation in the synchronized cells but were unsuccessful. Additionally, due to poor or unavailable antibodies for other DUSPs proteins we were limited to investigating the presented proteins but further evaluation of the remaining DUSPs is warranted.

Direct oxidation of ERK1/2 can also regulate its phosphorylation status. The oxidation of ERK1/2 upon the addition of serum is rapidly increased across all three cell lines but is sustained through the 2-4 hours in the Miro1^-/-^ MEFs, whereas it decreases in the Miro1^+/+^ and Myc-Miro1 MEFs. The oxidation of ERK1/2 in the Miro1^-/-^ MEFs could be the key factor in its dynamic regulation.

Overall, we observed a proliferative and cell cycle defect upon Miro1 deletion in MEFs which was accompanied by the first ever transcriptomic analysis dependent upon Miro1 expression. We have reason to believe that the oxidant rich perinuclear and nuclear spaces in the Miro1^-/-^ MEFs is leading to direct oxidation of ERK1/2 which could dysregulate its phosphorylation dynamics throughout the cell cycle and this defect is not due to the inactivity of ERK1/2 specific phosphatases. The involvement of ERK1/2 phosphorylation and the progression of the cell cycle dependent upon the expression of Miro1 has not been characterized before despite the mechanism not being fully revealed. Together, our studies provide strong support for global gene-expression changes and regulation of ERK1/2 dynamics via Miro1 expression. Data presented within will support investigating these signaling pathways identified in disease models in which Miro1 is implicated.

## Materials and Methods

### Cell culture

Miro1^+/+^ and Miro1^-/-^ mouse embryonic fibroblasts (MEFs) were provided by Dr. Janet Shaw and were cultured in antibiotic free 1x DMEM (Gibco) supplemented with 4.5g/L D-glucose, 10% fetal bovine serum (FBS, Gibco) and 78.9μM beta-mercaptoethanol (BMe, MP Biomedicals). Miro1^-/-^ Myc-Miro1 fibroblasts were generated as previously described [4]. Miro1^-/-^ Myc-Miro1 MEFs were cultured with 3mg/mL hygromycin b to select for cells with the Myc-Miro1 plasmid. These cells were grown in a humidified incubator at 37°C with 5% CO2.

### Cell proliferation

MEFs were plated at 25,000 cells/well of a 6-well plate with 4mL 10% FBS DMEM. Cells were trypsinized every 24 hours for 144 hours and counted using a hemocytometer and trypan blue staining. Media was changed every 48 hours. Cell counts were graphed as cells/mL versus time.

### Cell synchronization

Cells were grown to 70% confluency and treated with 0.1-1% FBS DMEM for 24-48 hours. Full serum media (10% FBS DMEM) was added back for the designated time points and cells were collected in lysis buffer (see Western blotting section) and quantified using a Bradford assay. 10-15μg of lysate were analyzed using a Western blot as described above.

### Cytosolic/nuclear fractionization

MEFs in a 6-well plate were scraped in 1mL ice-cold 1x phosphate buffered saline (PBS) and pelleted. The cell pellet was resuspended in 400uL ice-cold hypotonic buffer (10mM HEPES pH 7.9, 10mM KCl, 0.1mM EDTA, 0.1mM EGTA, 1mM DTT, 0.5mM PMSF, 10% IGEPAL) and left on ice for 5 minutes. Samples were centrifuged at 12,000 rpm for 30 seconds and the supernatant was collected as the cytosolic fraction. The pellet was washed twice in 800μL ice-cold 1x PBS and centrifuged at 10,000 rpm for 15 seconds. The pellet was then resuspended in 40μL ice-cold hypertonic buffer (20mM HEPES pH 7.9, 0.4M NaCl, 1mM EDTA, 1mM EGTA, 1mM DTT, and 1mM PMSF) and vigorously rocked at 4°C for 15 minutes. This was then centrifuged at 13,000 rpm for 15 minutes at 4°C and the supernatant was saved as the nuclear fraction.

### DCP-Bio1 labeling and Pulldown

To label sulfenic acids, the MEFs were lysed in lysis buffer (see Western blotting section). They were left on ice for 30 minutes then centrifuged at 10,000 x g for 5min at 4°C to pellet the debris. The supernatant was then passed through an Amicon filter (Millipore #UFC501096) and washed five times in washing buffer (50mM HEPES, 250mM NaCl, 1.5mM MgCl2, 1mM EDTA, pH7.4) to clear the unreacted DCP-Bio1. The lysate was resuspended in lysis buffer supplemented with protease inhibitor cocktail, 25mM NaF, and 2mM Na3VO4. Streptavidin magnetic beads (Promega #V7820) were added to 50μg lysate and allowed to rotate overnight at 4°C to pulldown the DCP-Bio1 labeled sulfenic acids. The magnetic beads were washed three times in lysis buffer (see Western blotting section) then the proteins were stripped from the beads using 5x sample buffer + 20% 1M DTT and heated at 95°C for 5 minutes. These samples were run on a 4-12% NuPAGE MOPs gel, as listed above, alongside 10μg lysate as a loading control.

### Flow cytometry

Cells were trypsinized, pelleted at 2,500 x g for 5 minutes, washed in 1mL 1x PBS then centrifuged at 3,000 x g for 5 minutes. Ice-cold 70% ethanol was added dropwise to the cell pellet while vortexing. These cells remained at 4°C for a minimum of 30 minutes then they were centrifuged at 3,000 x g for 5 minutes and washed in 1mL 1x PBS three times. The cell pellet was treated with 100μg/mL RNase (Sigma #10109142001) at room temperature for 5 minutes. After 5 minutes, 50μg/mL propidium iodide (Invitrogen #P21493) was added then cells were run through the Beckman CytoFlex with a 488nm laser, and the data was analyzed using FlowJo’s Watson (pragmatic) model.

### HyPer7 transfection and image analysis

For all three MEF cell lines, 200,000 cells were trypsinized and pelleted. The cell pellet was mixed with 500ng of plasmid (pCS2+MLS-HyPer7, Addgene # 136470) and taken up by the Neon Transfection System pipette. The cells were electroporated using the following conditions: 1350 volts, 30ms pulse width, and 1 pulse. The cells were grown on top of a glass coverslip overnight in 10% FBS DMEM. The next morning the cells were captured on an Eclipse TE-2000E inverted microscope (Nikon) equipped with a 40×/1.3 numerical aperture (NA) Plan Fluor oil-immersion objective. The MEFs were excited at 470nm and 440nm and the oxidation was calculated by dividing the fluorescence intensity at 470nm by the intensity at 440nm.

### Immunofluorescence staining and analysis

Cells were plated on glass coverslips at 100,000 cells per 6-well dish then fixed for 10 minutes with 4% paraformaldehyde/1x phosphate buffered saline (PBS), then permeabilized with 0.25% permeabilization buffer (triton 100-x in 1x PBS) for 10 minutes. The cells were then blocked in 1.5% BSA/1x PBS at 4°C overnight. The coverslips were incubated with TOMM-20 (1:250, Millipore #MABT166) for 1h, washed in 1x PBS for 40 minutes, incubated with Alexa Fluor 594 phalloidin (1:400, Invitrogen #A12381) and Alexa Fluor 488 (1:500, Invitrogen #A11008), washed in 1x PBS for 40 minutes, then stained with DAPI (0.5μg/mL, ThermoFisher #62248) for 5 minutes. Images were captured using an Eclipse TE-2000E inverted microscope (Nikon) equipped with a 40×/1.3 numerical aperture (NA) Plan Fluor oil-immersion objective. Mitochondrial occupancy was calculated by drawing the area of the mitochondria in the leading and lagging edges of the cell and that was taken as a percentage of the occupancy in relation to phalloidin in those areas. Tomm-20 fluorescence intensity was done by highlighting the mitochondria and taking the average fluorescence intensity from each cell. The mitochondrial form factor was calculated by highlighting the mitochondria and taking their area and perimeter and plugging it into the equation: Form Factor = 0.25 (area/perimeter^2^).

### Quantitative real-time PCR

Cells were pelleted and then frozen in liquid nitrogen until the RNA was extracted using the RNeasy kit (Qiagen #74104) according to the manufacturer’s protocol. The RNA was converted to cDNA using the RT^2^ First Strand Kit (Qiagen #330404) and then analyzed for the following primers: cdh2 (IDT #Mm.PT.58.12378183) and beta-actin (IDT #Mm.PT.58.33540333). The primers, cDNA, and RT^2^ SYBR Green ROX (Qiagen #330523) were mixed into a 96-well plate and run on the Quant Studio 3 instrument.

### RNAseq and analysis

Biological replicates of RNA isolated from all MEF lines were submitted to Novogene for QC analysis, RNA library prep and paired-end sequencing. Subsequent FASTQ files were aligned with STAR (v2.7.10b) using the mouse GRCm39 assembly and gencode vM34 annotation. Read counts were normalized and differential gene expression performed with DESeq2 (v1.44.0). Pathway enrichment analysis was performed on DEGs identified between Miro1^+/+^ and Miro1^-/-^ MEFs using R (v4.4.0) and the PathfindR package (v2.4.1). Heatmaps were created using Pheatmap (v1.0.12) and volcano plots with ggplot2 (v3.5.1). All custom scripts used for figure generation will be made available upon request.

### Western blotting

Cells were collected in lysis buffer (50mM HEPES, 250mM NaCl, 1.5mM MgCl2, 1mM EDTA, 10% Glycerol, 1% Triton-X100, 25mM NaF, and 2mM Na3VO4 pH 7.4) supplemented with protease inhibitor (Pierce #A32955). The protein concentration was quantified using a Bradford assay. Then took 10-15μg protein and mixed with sample buffer to a final concentration of 1x sample buffer (1M Tris pH6.8, 10% SDS, 100% Glycerol, 3mg Bromophenol blue) supplemented with 20% 1M dithiothreitol (DTT), was loaded per lane on a 4-12% NuPAGE MOPs gel which was run at constant 200 volts for 50 minutes in the presence of 1x MOPs buffer. The protein was transferred to a PVDF membrane from Cytiva Amersham (#10600023) in the presence of 1x transfer buffer (150mL NuPAGE transfer buffer (20x), 600mL methanol, 2.25L dH2O). The transfer was done at 4°C for 90 minutes at constant 0.50 Amps. Membranes were then blocked in 5% bovine serum albumin (BSA)/1x tris buffered saline (TBS) with 0.2% Tween-20 (TBST) at room temperature for 1h. The membranes were immunoblotted at 4°C overnight with the following antibodies in 5% BSA/1x TBST: MKP1/DUSP1 (1:1000, Invitrogen #MA5-32480), DUSP3 (1:2000, Abcam #AB125077), DUSP5 (1:2000, Origene #CF11035) MKP3/DUSP6 (1:1000, Invitrogen #MA5-35048), PP2AC (1:1000, Cell Signaling Technology #2259), GAPDH (1:3000, Invitrogen #MA5-15738), Cyclin D1 (1:1000, Cell Signaling Technology #2978), Cyclin B1 (1:1000, Cell Signaling Technology #4138), Cyclin A2 (1:1000, Abcam #AB32386), phospho-ERK1/2 (T202/Y204) (1:1000,

Cell Signaling Technology #9101), ERK1/2 (1:2000, Cell Signaling Technology #9102), phospho-MEK1/2 (S217/S221) (1:1000, Cell Signaling Technology #9121), total MEK1/2 (1:1000, Cell Signaling Technology #9122), Cofilin (1:1000, Abcam #AB42824), Lamin A/C (1:1000, BD Transduction Laboratories #612162), cofilin (1:1000, Abcam #AB-42824), Tomm-20 (1:1000, Millipore #MABT166) The blots were then imaged using the Amersham Imager 600.

### Statistical analysis

Statistics were conducted using GraphPad Prism 10.2.3. Statistical tests used are noted in the figure legends.

## Acknowledgement

We thank Dr. Alan Howe, University of Vermont Department of Pharmacology for generous use of his microscope. We thank the Harry Hood Bassett Flow Cytometry and Small Particle Detection Facility (RRID:SCR_022147) at the University of Vermont Larner College of Medicine for flow cytometry support.

