## Supplementary figures and images for "Miro1 expression alters global gene expression, ERK1/2 phosphorylation, oxidation, and cell cycle progression"

### Supplemental Figure 1

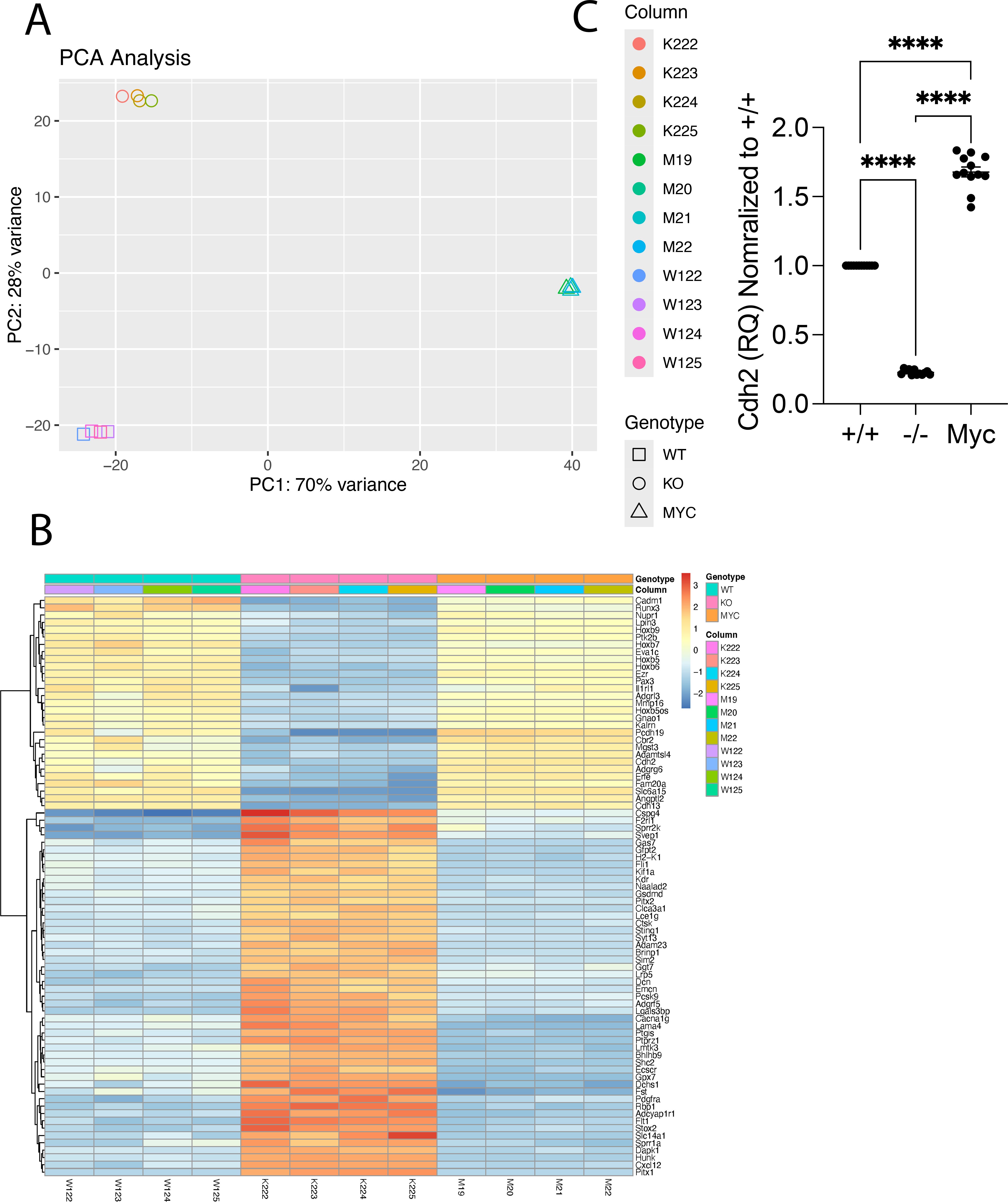
